# Molecular mechanisms of sperm motility are conserved in a basal metazoan

**DOI:** 10.1101/2021.05.28.446218

**Authors:** Kelsey F. Speer, Luella Allen-Waller, Dana R. Novikov, Katie L. Barott

## Abstract

Efficient and targeted sperm motility is essential for animal reproductive success. Studies in mammals and echinoderms have uncovered a highly conserved signaling mechanism in which sperm motility is stimulated by pH-dependent activation of the cAMP-producing enzyme soluble adenylyl cyclase (sAC). However, the presence of this pathway in basal metazoans has, until now, been unexplored. Here we found that cytoplasmic alkalinization induced a rapid burst of cAMP signaling and the full activation of motility in sperm from the reef-building coral *Montipora capitata*. Coral sperm expressed sAC in the flagellum, midpiece, and acrosomal regions, indicating that this molecular pH sensor may play a role in regulating mitochondrial respiration and flagellar beating. In bilaterians, sAC is a central node of a broader pH-dependent signaling pathway that alters cellular behavior in response to changes to the extracellular environment. We present transcript-level evidence that a homologous pathway is present in coral sperm, including the Na^+^/H^+^ exchanger SLC9C1, protein kinase A, and the CatSper Ca^2+^ channel conserved even in mammalian sperm. Our discovery of this pathway in a basal metazoan species highlights the ancient origin of the pH-sAC-cAMP signaling node in sperm physiology and suggests that it may be present in many other marine invertebrate taxa for which sperm motility mechanisms remain unexplored. These results emphasize our need to better understand the role of pH-dependent signaling in marine reproductive success, particularly as worsening ocean acidification and warming due to climate change continue to impair the physiology of corals and other marine invertebrates.

**Statement of significance:** Reef-building corals are the keystone species of the world’s most biodiverse yet threatened marine ecosystems. Corals reproduce by broadcast spawning, making the ability of their sperm to swim through the water column essential for fertilization. However, little is known about the mechanisms that regulate coral sperm motility. Here we found that elevated intracellular pH promotes the production of the second messenger cAMP in coral sperm and triggers the onset of motility. This study reveals the deep conservation of a sperm activation pathway from humans to corals, presenting the first comprehensive examination of the molecular mechanisms regulating sperm motility in an ancestral animal. These results are critical for understanding the resilience of this sensitive life stage to a changing marine environment.

## Introduction

Activation of sperm motility requires precise temporal and spatial control in order to maximize chances of egg-sperm contact and fertilization (1). Despite the vital importance of this process for reproductive success across metazoan phyla, the molecular mechanisms that regulate sperm activation remain poorly understood. Sperm are typically held in an inactive state within the male prior to release, either into the surrounding water column during broadcast spawning, as in basal metazoans and many bilaterians, or directly into the female oviduct during copulation, as in some bilaterians (2). The signals that trigger sperm motility following release vary between environments and species, and can include changes in osmolarity, ion concentrations (e.g. bicarbonate), and/or chemical signals released by eggs (3). Even with the considerable differences in reproductive ecology and activation cues that characterize different taxa, the downstream signaling pathways that activate motility are highly conserved across the few taxa that have been described to date, namely mammals (phylum Chordata) and sea urchins (phylum Echinodermata). Because of the historical focus on these two phyla, which are closely related in the context of metazoan evolution, we know very little about the broader evolution of the molecular pathways that activate sperm motility across the metazoan phylogeny, and especially in more basal metazoan phyla. This has left a significant gap in our understanding of the evolution of the mechanisms that regulate sperm function, an important issue to address given the importance of these mechanisms for determining animal fitness in a changing environment.

Reef-building corals, members of the basal phylum Cnidaria, are an evolutionarily and ecologically important model system for understanding mechanisms of sperm motility. First, corals belong to one of the earliest phyla to evolve organized tissues, and as diploblasts, they lack the mesoderm present in bilaterians (4). Second, corals are the foundational species of coral reefs, one of the most biodiverse ecosystems on the planet (5). Coral reproduction is essential for the persistence of reefs worldwide (6), and in most species involves the tightly coordinated release of sperm into the water column on just a few nights each year (7, 8). This process is currently threatened by a combination of local stressors (9) and global climate change (10–12). Sperm, in particular, are an acutely susceptible life stage to environmental disturbances due to their small size and brief life span (13), and climate change stressors including ocean warming and acidification have reduced coral sperm production (11, 14), motility (10, 15, 16) and fertilization success (17, 18). The mechanisms driving these declines in sperm performance are unknown, but both warming and acidification may disrupt coral cellular metabolism and acidify the cytosol (19–21), two processes that are important for sperm motility in other species (22). Indeed, initial evidence indicates that alkalinization of coral sperm cytosol promotes motility (23), highlighting the importance of understanding the molecular pathways that connect pH-dependent signaling with changes in cellular performance.

Alkalinization of the sperm cytosol acts as a critical intracellular messenger controlling the onset of motility in several species (3, 24, 25). In sea urchins, binding of egg-derived peptides (e.g., speract; (2)) to guanylyl cyclase (GC) receptors at the cell surface activates a sperm-specific Na^+^/H^+^ exchanger (SLC9C1; (26)), which increases cytoplasmic pH through its proton efflux activity. In both spawning marine invertebrates and mammals, this alkalinization stimulates cAMP production via the enzyme soluble adenylyl cyclase (sAC; (27))), leading to protein kinase A (PKA)-dependent phosphorylation of flagellar proteins and calcium signaling via CatSper channels (2), which together stimulate flagellar beating. Cnidarian GC-A receptors and CatSper channels are highly conserved with those from sea urchins (28, 29), and although the other constituents of the pathway have yet to be investigated, these reports suggest that this molecular mechanism may be conserved in basal metazoans. Mammals have evolved a distinct activation signal that nonetheless utilizes a similar signaling cascade to that of sea urchins, whereby elevated levels of bicarbonate in the female reproductive tract activate sAC (30), and the resulting burst of cAMP activates PKA, leading to phosphorylation of flagellar proteins and ultimately motility. Each component of this conserved pathway appears to be necessary for sperm motility and fertilization in mammals, as mice lacking SLC9C1, sAC, PKA or CatSper display severe sperm motility defects, rendering them infertile (31–35). These studies support the conservation of sAC-cAMP as a central signaling node in sperm activation across bilateria, however, there is no data currently on the role of sAC-cAMP signaling in basal metazoans. In corals, somatic tissues express a functional homolog of sAC that is stimulated by bicarbonate to make cAMP (36), and this enzyme plays a role in responding to pH fluctuations within the cell (37), leaving open the promising possibility that this pathway is functionally conserved in coral sperm.

In order to test the hypothesis that sperm motility in basal metazoans is regulated by a molecular signaling pathway that is conserved with bilaterians, we examined the role of intracellular pH, sAC and cAMP signaling in sperm motility in the reef-building coral *Montipora capitata* using a combination of microscopy and immunological and biochemical assays. In addition, we analyzed the expression and conservation of key proteins in the echinoderm motility initiation pathway in *M. capitata* sperm via interrogation of RNA-seq databases and *in silico* structural analyses. This is the first comprehensive examination of the intracellular signaling pathways that regulate sperm motility in a basal metazoan, and our results highlight the functional conservation of a fundamental pathway essential for animal reproduction. Furthermore, the mechanisms underlying coral sperm motility have important implications for determining how climate change will influence the reproductive success of corals and other marine animals.

## Results

### Response of coral sperm to cytosolic alkalinization

Sperm from the coral *Montipora capitata* were exposed to 20 mM NH_4_Cl to induce cytosolic alkalinization, and intracellular pH (pH_i_) and motility were monitored simultaneously by confocal microscopy using the fluorescent dye SNARF-1-AM (Fig. S1). Prior to NH_4_Cl treatment, sperm were inactive (0% motility) and had a mean pH_i_ of 7.52 ± 0.02 (Fig. 1A). Sperm pH_i_ increased following NH_4_Cl exposure, reaching a peak of 8.31 ± 0.04 within 45 seconds (Fig. 1A), equivalent to over an 80% decrease in H^+^ concentration. Sperm pH_i_ remained above pH 8.0 for at least 2:30 min, then started to decline after 3:00 min, reaching initial levels after 4:30 min. pH_i_ continued to decline over the next 2:00 min, reaching a minimum of pH 7.26 ± 0.02 (Fig. 1A). Motility increased instantaneously following exposure to NH_4_Cl (visual observation), reaching 100% within 45 seconds post-exposure (Fig. 1A). Motility remained near 100% for 3:20 min, then rapidly decreased to 7.1% after 4:25 min and remained near zero until the end of the experiment (Fig. 1A). This drop in motility was concurrent with declining pH_i_, with a possible motility threshold of pH_i_ ~7.8. Exposure of sperm to 20 mM NH_4_Cl also led to a 13.5-fold increase in cAMP content, from 0.129 ± 0.009 nmol cAMP mg^−^ protein to 1.747 ± 0.248 nmol cAMP mg^−^ protein within 5 seconds (Fig. 1B). This burst of cAMP was transient, declining by 58% within 20 seconds of the peak and by 78% at 60 seconds post-exposure. Sperm cAMP concentrations remained 2-3-fold higher than initial levels for up to 5 min post-exposure while unexcited control sperm exhibited no change in cAMP content (Fig. 1B).

**Figure 1.**
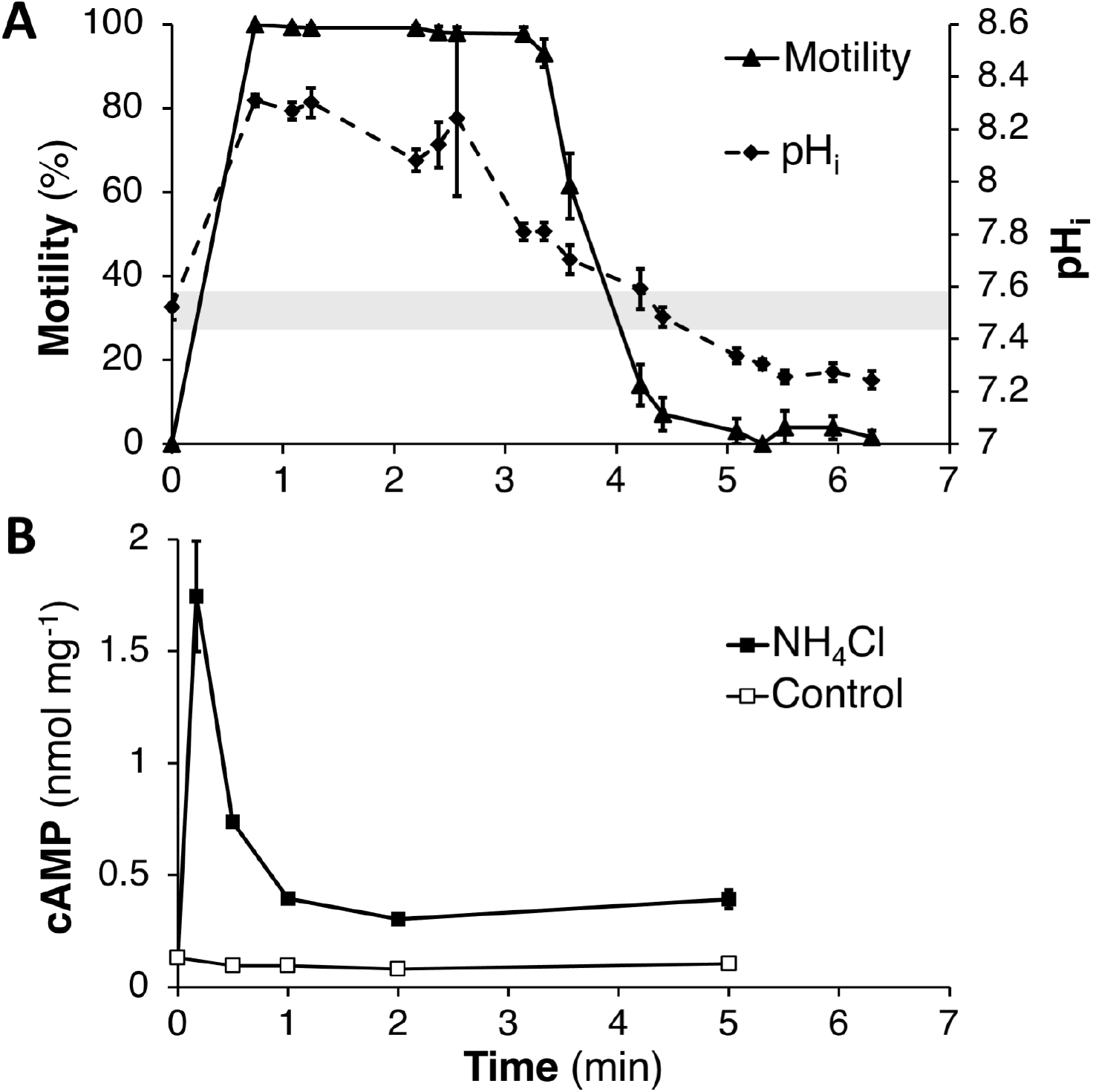
Response of sperm from the coral *Montipora capitata* to ammonium chloride (NH_4_Cl) treatment. A) Coral sperm motility (solid line) and pH_i_ (dashed line) after exposure to 20 mM NH_4_Cl (N ≥ 13 cells per time point). Gray shaded region indicates 99% confidence interval of sperm inital pH_i_ before activation (7.53 +/− 0.06). B) Concentration of cAMP in coral sperm, normalized to total protein, following treatment with 20 mM NH_4_Cl (black squares) or control (open squares); N=3. All error bars indicate SEM; where not visible they fall within the symbol.

### Expression of soluble adenylyl cyclase in coral sperm

The sequence of *M. capitata* sAC (*mc*sAC) was identified by querying an *M. capitata* sperm genome (38) using a cDNA sequence from the coral *Pocillopora damicornis* (*pd*sAC; (37)). The predicted protein was 75.7% similar to *pd*sAC, and 48.3% similar to sea urchin sAC (Table S1). Sequence alignment with metazoan homologs (Table S2) demonstrated that *mc*sAC contained both the conserved adenylyl cyclase catalytic domains necessary for bicarbonate-stimulated cAMP production (39), and a long (~140 kD) C-tail of unknown function. Like *pds*sAC, *mc*sAC lacked the short autoinhibitory peptide directly C-terminal to catalytic domain 2 (Fig. S2), which has been described in mammals (40). Expression of *mc*sAC in sperm was confirmed by Western blot using anti-coral sAC antibodies against the second catalytic domain, which indicated that the predominant isoform was the full-length protein (~195 kD; sAC_FL_; Fig. 2A,B), followed by a ~110 kD isoform (sAC_110_, Sperm (high); Fig. 2A), and two additional isoforms detected at low levels that were ~55 kD and 45 kD (sAC_55_ and sAC_45_, respectively; Sperm (low), Fig. 2A). Expression of sAC_FL_ was not detected in adult *M. capitata* tissues, which only expressed the smallest ~45 kD isoform at detectable levels, which ran as a doublet (Fig. 2C), and may represent post-translational modifications of sAC_T_ or possibly an atypical variant of sAC lacking CY1 found in mammalian somatic cells (41).

**Figure 2.**
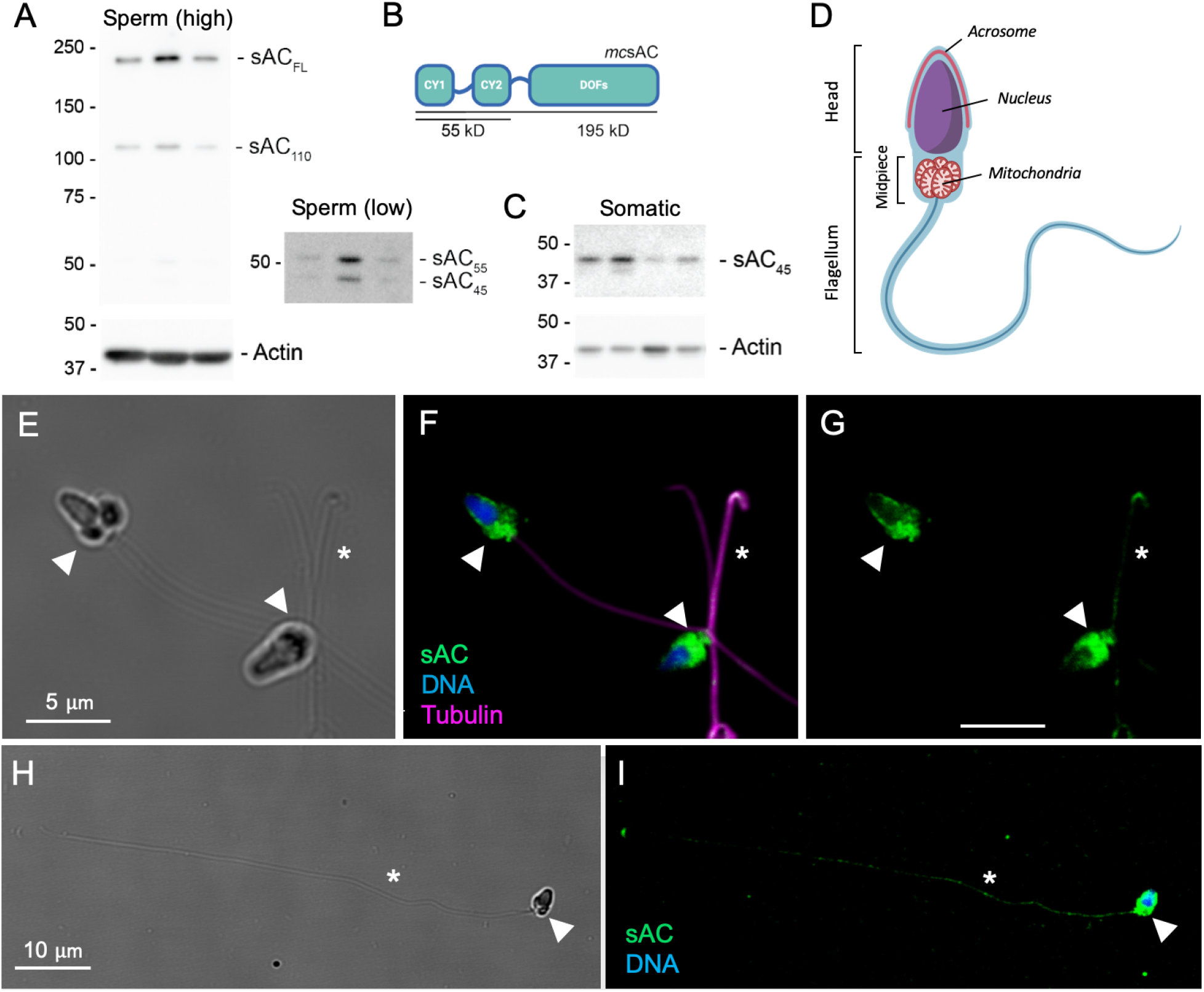
Expression of sAC in *Montipora capitata* sperm. Western blots of the sAC protein show A) high expression of two isoforms (Sperm (high); sAC_FL_, sAC_110_) and low expression of two isoforms (Sperm (low); sAC_55_, sAC_45_) in sperm from three different individuals. Numbers indicate approximate protein size in kD. B) *In silico* structural analysis predicts that *M. capitata* sAC (*mc*sAC) contains two catalytic cyclase domains (CY1 & CY2; ~55kD) followed by several domains of unknown function (DOFs) that, in total, encode a protein of ~195 kD. C) Adult *M. capitata* somatic tissues expressed a single sAC isoform (sAC_45_). D) Diagram of the architecture of coral sperm, identified by (42). The head houses the nuclei and the surrounding acrosome compartment. The flagellum contains the axoneme formed from a 9 + 2 microtubule bundle and, immediately posterior to the head, a mitochondria-rich midpiece. E) Brightfield image of sperm; F) Corresponding fluorescence micrograph showing localization of sAC (green), DNA (blue), and flagella (magenta; stained with anti-β-tubulin antibodies); G) Corresponding image of coral sAC expression alone. H) Brightfield image of a sperm cell with the flagellum extended; I) Corresponding fluorescence micrograph highlighting coral sAC expression along the length of the flagellum. Arrowheads indicate the midpiece of the flagellum containing the mitochondrial sheath and asterisks indicate the flagellum.

The subcellular localization of *mc*sAC in sperm was determined by immunocytochemistry using the same anti-coral sAC antibodies. *M. capitata* sperm consist of a head that contains the nucleus and acrosomal compartment, and a flagellum with a single 9 + 2 microtubule bundle that includes two distinct regions: the midpiece adjacent to the head that contains 5-6 mitochondria, followed by the principle piece (i.e. tail), containing just the axonemal fibers (Fig. 2D; (42)). *M. capitata* sperm expressed *mc*sAC throughout the entire cell, and expression was most concentrated in the midpiece (Fig. 2E-G). *mc*sAC was also present in low abundance at the tip of the sperm head in the predicted acrosomal region (Fig. 2E-G) and across the entire length of the flagellum (Fig. 2H, I), and it co-localized with β-tubulin, the primary component of the axonemal fibers (Fig. 2F, G). Controls confirmed antibody staining was specific for *mc*sAC for both Westerns (Fig. S3B, D) and immunostaining (Fig. S4).

### Expression of a conserved bilaterian signaling core in coral

Next we next analyzed the expression of key players in the sAC-dependent sperm activation pathway in coral sperm; both those conserved across bilateria (PKA, SLC9C1 [sNHE], CatSper) and those that have only been described in marine invertebrates (GC-A, HCN, and CNGK) (Fig. 3A). We began by searching in two *M. capitata* genomes (38, 43) and a sperm RNA-seq database (44) using sea urchin homologs. Further structural analysis was carried out using sequence alignment and transmembrane helix prediction software in order to identify key functional domains (Table S2 & S3; Fig. 3C-G; Fig S5-8). In cases where the *M. capitata* genome predicted a structurally incomplete protein, a comparison with the well-annotated genome of *P. damicornis* was used to identify potential missing sequences (dotted lines; Fig. 3C-G). Together these analyses identified a single homolog of each gene in the *M. capitata* genome and at least 9 transcripts of 95% or greater homology to the genomic sequence in the sperm transcriptome (Table S1). Specifically, the catalytic domain of PKA in *M. capitata* sperm was 90.6% similar to *S. purpuratus* PKA Cα, encoding a polypeptide of ~40 kD (Table S1). Protein expression of PKA Cα in *M. capitata* sperm was further confirmed by Western blot (Fig. 3B). The band ran as a triplet, likely due to activating phosphorylation of the kinase activation loop (45). The *M. capitata* homolog of SLC9C1 (i.e. sNHE) was 69.8 % similar to SLC9C1 from *S. purpuratus* (Table S1). *mc*SLC9C1 contained an N-terminal sodium-proton exchanger (NHE) domain with the canonical cation-binding motif of 1:1 electroneutral exchangers embedded within the predicted S3-4 region, identical to echinoderms but distinct from mammals (Fig. 3C; Fig. S5A). C-terminal to the NHE, *mc*SLC9C1 also had both a voltage-sensing domain (VSD) and a cyclic nucleotide-binding domain (CNBD; Fig. 3C). The *mc*SLC9C1 VSD displayed the four-transmembrane (S1-4) architecture, including the seven positively charged residues in S4, found in the *Drosophila* Shaker channel (Fig. S5B; Aggarwal and MacKinnon 1996). While these charged residues are entirely conserved in echinoderms, they are lost in mammals, reflecting the divergence of the pathway from its invertebrate ancestors (Wang et al., 2003).

**Figure 3.**
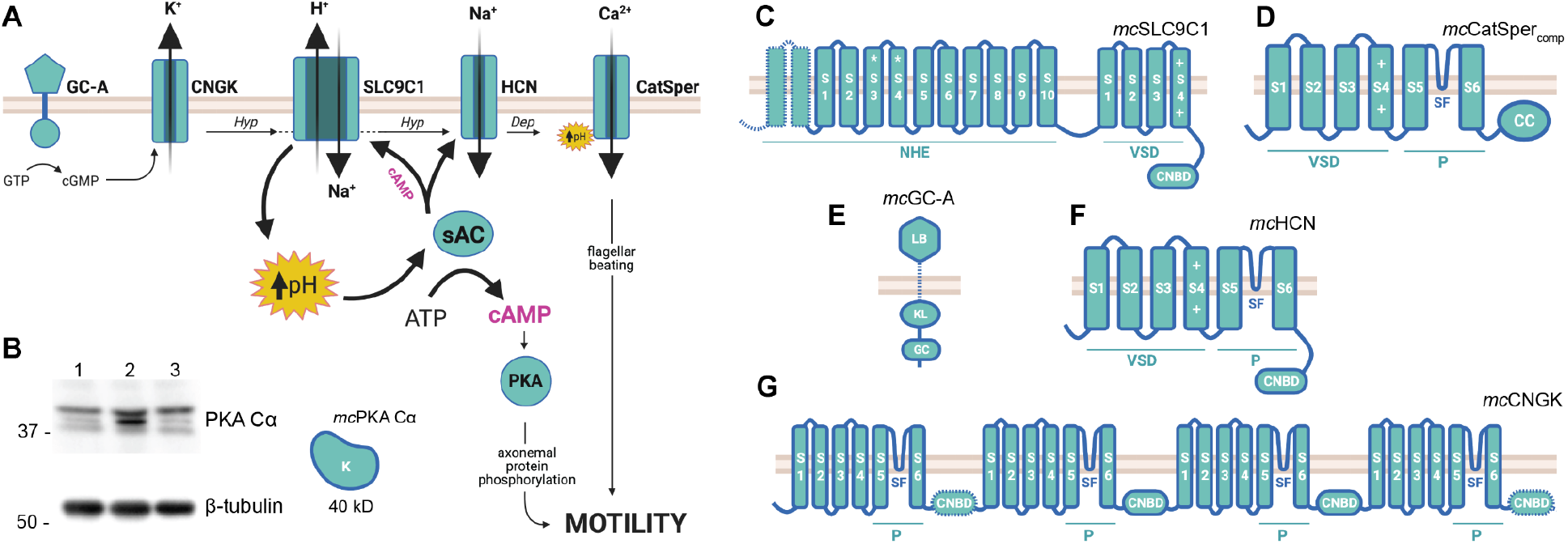
A pH-dependent motility pathway is conserved in sperm from the coral *Montipora capitata*. A) Diagram of the pH-sAC-cAMP motility pathway from echinoderms. Egg-derived chemoattractants bind to a guanylyl cyclase receptor (GC-A) which produces cGMP to stimulate CNGK-mediated K^+^ efflux and membrane hyperpolarization (Hyp). CNGK activates the H^+^/Na^+^ exchanger SLC9C1 which raises cytoplasmic pH, thereby activating sAC-dependent cAMP production and driving PKA-dependent phosphorylation of axonemal proteins controlling motility. sAC-dependent cAMP feeds back to maintain SLC9C1 activity and promotes hyperpolarization-dependent activation of HCN, which depolarizes (Dep) the cell via Na^+^ influx. The CatSper channel responds to both depolarization and elevated pH to generate Ca^2+^ influx signals that alter the flagellar waveform. B) Western blot analysis of PKA Cα expression in *M. capitata* sperm yielded three bands close to the predicted protein size of 40 kD. Predicted domain structures of *M. capitata* homologs of C) *mc*SLC9C1; D) *mc*CatSper_comp_; E) *mc*GC-A; F) *mc*HCN; and G) *mc*CNGK. Conserved protein domains and amino acid signatures: K, kinase domain; S, transmembrane segment; NHE, sodium hydrogen exchange domain; *, includes NHE consensus sequence; VSD, voltage sensing domain; +, positively charged amino acids involved in voltage sensing; CNBD, cyclic nucleotide-binding domain; P, pore-forming domain; SF, selectivity filter; CC, coiled-coil domain; CY, cyclase domain; KL, kinase-like domain; LB, ligand-binding domain.

The four polypeptides that assemble to form the pore of the CatSper calcium ion channel (CatSper1-4ɑ; (46)) were identified in *M. capitata* sperm and shared up to 70.7% similarity with their *S. purpuratus* homologs, but were 27.7 - 74.3% shorter, likely due to incomplete genome coverage (Table S1; Fig. S6A). When the partial sequences were overlaid, a consensus of the classic 6-TM type architecture common to CatSpers emerged (Fig. 3D). For example, *mc*CatSper2-4ɑ each contained a VSD with 4-5 charged amino acids in transmembrane segment S4, as compared with the 6-7 found in the sea urchin *A. punctulata* and the 2-4 found in *Homo sapiens* (Fig. S6B). In *mc*CatSper1, 2, and 4ɑ, segments S5 and S6 were linked by a short interconnecting helix, or “selectivity filter” with the canonical [T/S]x[D/E]xW motif of Ca^2+^-selective channels (Fig. 3D; Fig. S6C). *mc*CatSpers 2 & 3ɑ contained short, cytoplasmic domains C-terminal to S6 that were predicted to form coiled-coil domains, important for oligomerization in mammals (Table S4; Fig. S6A; (46, 47)). Transcripts of CatSper2-4ɑ were identified in *M. capitata* sperm (Table S1), and while *mc*CatSper1ɑ was not found, it is notable that the stable expression of the four bilaterian subunits is highly interdependent (48). Thus the absence of *mc*CatSper1ɑ is likely due to an inability to identify the C-terminus of the protein via homology searches of the genome, which combined with the inherent 3’ bias of the poly-A selection method used to generate the RNA-seq library (44), likely obscured actual expression data.

Considering the invertebrate-specific members of the pathway, the *M. capitata* guanylyl cyclase receptor (*mc*GC-A) gene was 42.3% similar to *S. purpuratus* GC-A (synonymous with the Speract receptor) and 78.0% similar to that of the coral *Euphyllia ancora* GC-A (*ea*GC-A; Table S1). *mc*GC-A contained an intracellular kinase-like homology (KL) domain and a guanylyl cyclase catalytic domain (CY), as well as an extracellular ligand-binding domain (LB; Fig. 3E), similar to both the Speract receptor and *ea*GC-A. Sea urchin sperm activation via GC-A also involves the ion channels CNGK and HCN. CNGK-dependent membrane hyperpolarization activates SLC9C1 to induce cytosolic alkalinization, whereas HCN-dependent membrane depolarization occurs downstream of sAC to facilitate CatSper activation (Fig. 3A; (2)). Like CatSper, both HCN and CNGK assemble as a tetramer of 6TM-type subunits, however with a C-terminal CNBD replacing the CatSper coiled-coil domain (Fig. 3F; Fig. 3G, respectively). *mc*CNGK was 53.9% similar to CNGK characterized in the sea urchin *Arabica punctulata* (Table S1), and its four subunits were encoded as domains within a single polypeptide (Fig. 3G). In contrast, HCN channels form a homotetramer (49), and the single 6-TM repeat of *mc*HCN (Fig. 3F) was 73.1% similar to the *S. purpuratus* homolog (Table S1). Like their echinoderm homologs, the selectivity filters of both *mc*CNGK and *mc*HCN contained a canonical cation-binding GYG motif of cation channels (Fig. S7; Fig. S8A). This region in *mc*HCN was nearly identical to *sp*HCN (100% similarity; Fig. S8A), a channel whose weak selectivity for K^+^ (P_K_/P_Na_ = 4.7; (50)) causes a depolarizing inward Na^+^current under physiological conditions (51). In contrast, the T/S rich region in the *mc*CNGK filter (Fig. S7) is indicative of highly K^+^-selective channels (51). Additionally, *mc*HCN transmembrane segment S4 contained eight positively charged amino acids, broken into two clusters of 3-4 residues, that comprise a predicted VSD (Fig. S8B). *mc*CNGK, however, lacked a VSD, suggesting this channel responds solely to cGMP and not to changes in membrane polarization.

## Discussion

### Cyclic AMP is a conserved signaling mechanism in sperm activation

Here we demonstrate for the first time in a basal metazoan that sperm motility is initiated via a pH and cAMP-dependent signaling pathway. Chemical alkalinization of inactive *Montipora capitata* sperm, which reached a peak of pH 8.2, was sufficient to induce 100% motility. A subsequent reacidification of the cytosol began ~2.5 minutes after alkalinization, and intracellular pH (pH_i_) passed below the initial pH of 7.5 to reach a minimum of pH 7.2. This overcompensation is consistent with the presence of active acid-base compensatory mechanisms in coral sperm. Once pH_i_ declined below pH 7.8, motility also began to decline, indicating that coral sperm may have a threshold for motility around pH_i_ 7.8, similar to observations in echinoderm sperm (24). While the mechanistic link between declining pH_i_ and reductions in motility in coral sperm remain uncharacterized, data from other marine invertebrate models suggests that the dynein ATPase, the key motor protein driving flagellar beating, is most active when pH_i_ is ≥ 7.5, and may be directly inhibited by reacidification (22, 52). Cytosolic alkalinization in coral sperm also coincided with rapid cAMP production that peaked within the first 5 seconds of activation, indicating that this universal second messenger molecule is involved in the onset of coral sperm motility. A similar burst of total cAMP occurs in both mammalian (53) and echinoderm sperm (54), and in those taxa initiates a signaling cascade that upregulates flagellar beating and alters the flagellar waveform (2). Our data suggest that this pH-dependent cAMP signaling is not unique to bilaterian sperm, but instead arose before the split of the cnidarian and bilaterian lineages.

### Soluble adenylyl cyclase may play a central role in coral sperm physiology

Here we described the expression and localization of the molecular pH sensor soluble adenylyl cyclase (sAC) for the first time in sperm from a basal metazoan. In bilaterians, sAC is the primary source of cAMP driving the onset of sperm motility, and plays a central role in the regulation of sperm activity (30, 55). Coral sAC activity is functionally conserved, exhibiting the hallmark activation by bicarbonate to generate cAMP (36). While the role of transmembrane adenylyl cyclases (tmAC) in coral sperm cAMP production cannot be ruled out, these enzymes are insensitive to pH and bicarbonate and do not play a major role in flagellar motility in bilaterian sperm (27, 56, 57). In contrast, sAC is activated in response to local fluctuations in sperm cytoplasmic pH in both mammals (58) and echinoderms (59), and is essential for motility (33). In coral sperm, sAC was expressed throughout the head, and in both the midpiece and distal region of the flagellum, indicating that sAC may be involved in multiple aspects of coral sperm physiology. For example, abundant expression along the midpiece, a localization that is conserved in sea urchin (59) and mammalian sperm (30), suggests that coral sAC may influence cellular respiration. In sperm, motility and respiration are tightly coupled through the dynein ATPase, which consumes the vast majority of cellular ATP (22). Indeed, studies of bilaterian somatic cells have uncovered a unique pool of sAC that resides within the mitochondrial matrix and promotes PKA-dependent upregulation of oxidative phosphorylation (60). Although the influence of sAC activity on sperm mitochondrial function remains unexplored, it is possible that regulation of mitochondrial output via sAC is an additional means of “tuning” flagellar motility. Importantly, because mitochondrial function and sperm motility are both impaired under simulated ocean acidification in sea urchin, mussel, and ascidian sperm *(21, 61)*, a description of the molecular mechanisms underlying sperm motility and their response to environmental conditions are critical for predicting the susceptibility of reproduction to climate change.

Localization of coral sAC along the entire length of the flagellum suggests an additional specialized role for this enzyme in motility. sAC also associates closely with the axoneme in sea urchins (59), where the large surface-area-to-volume ratio allows for rapid transmembrane signalling and direct contact with axonemal proteins (62). In addition, the other components of the motility pathway all localize almost exclusively to the sea urchin flagellum, including GC-A and CNGK (63), SLC9C1 (26), HCN (51), and CatSper (64), allowing for tight coupling of this signal transduction pathway. Although it remains to be determined where these proteins are expressed in coral sperm, it is likely that they similarly colocalize with coral sAC along the flagellum. In mouse sperm, sAC is expressed primarily in the midpiece but is not detected by immunocytochemistry in the principle piece (30). However, sAC activity has been observed in both of these compartments in mice, albeit with different kinetics (56), and the molecular mechanisms driving these differences remain to be described. Finally, the expression of coral sAC in the sperm head suggests it may play a role in the acrosome reaction, much like it does in sea urchins (59) and possibly in mammals (55). Interestingly, sea urchin sperm exhibit compartment-specific differences in sAC isoform expression, with sAC_T_ localizing to the head and sAC_FL_ along the flagellum (59). The functional significance of this partitioning remains unknown, but could lead to differences in enzyme kinetics between the head and the flagellum that influence their disparate roles in the acrosome reaction and motility, respectively.

Expression of multiple sAC isoforms due to alternative splicing is common in metazoans (65), and corals are no exception (Fig. 2; (37)). Coral sperm expressed multiple isoforms of sAC, including the first observation to our knowledge of sAC_FL_ expression in any cnidarian. That coral sAC_FL_ expression was restricted to the male germline, as it is in both mammals (30, 39) and echinoderms (59), suggests that there may be a conserved sperm-specific function of this isoform. However, the precise role of sAC_FL_ in cellular physiology remains enigmatic across all species examined to date. Interestingly, the autoinhibitory peptide described in mammalian sAC_FL_ (40) is absent in *mc*sAC and in sAC from all other coral species examined so far, which may contribute to the relatively high AC activity observed in corals (36) and highlights the need to understand more about the role of the sAC C-tail across metazoan species. Coral sperm also expressed several shorter isoforms, including a ~100 kD sAC isoform and two other small ~45-55 kD isoforms in low abundance. These smallest isoforms were also expressed in somatic coral tissues and may represent sAC_T_, an enzymatically active isoform containing only the two N-terminal cyclase domains, which is the minimal sequence required for bicarbonate-stimulated AC activity. sAC_T_ is commonly expressed in both germline and somatic cells of bilaterians (39, 59), agreeing with what we observed here in corals. All together, these data suggest that sAC plays a conserved and complex role as a central regulator of sperm physiology across metazoa, and much work remains to unravel the many potential roles of this enzyme in sperm physiology of corals and other basal metazoans.

### Conservation of a cAMP-dependent motility pathway across metazoans

A comprehensive analysis of the molecular mechanisms underlying sperm activation in coral confirmed the expression and high structural conservation of the entire echinoderm sperm activation pathway in a basal metazoan. These results indicate that coral sperm activation couples environmental sensing to a cytoplasmic signaling pathway dependent on intracellular alkalinization and the central regulatory node of sAC-cAMP. The ability of sperm to remain in a quiescent state until they detect an egg nearby allows males to optimize their fertilizing capacity by saving their limited energy and increasing their chances of encountering eggs by using directional motility towards the cue (i.e. chemotaxis) (2). The majority of extant spawning marine invertebrates utilize chemotaxis, including corals, various other cnidarians, molluscs, echinoderms, and ascidians (66), and this capacity is critical for fertilization (Fig. 4). In each of these lineages, we find functional data supporting the conservation of the pH-sAC-cAMP pathway (Fig. 4; (2, 67)). Genetic evidence for the co-evolution of sAC, SLC9C1, and CatSper also links this pathway to even more basal phyla, including Ctenophora and Porifera (Fig. 4; (68)). As copulation evolved in multiple lineages, this molecular mechanism was either lost (i.e., Nematoda) or it took on new roles, such as the sAC-dependent sperm maturation process in mammals (i.e., capacitation) (Fig. 4; (30)). Characterizing the evolution of this pathway will allow us to better understand both the shared and divergent traits underlying male fertility in metazoans.

**Figure 4.**
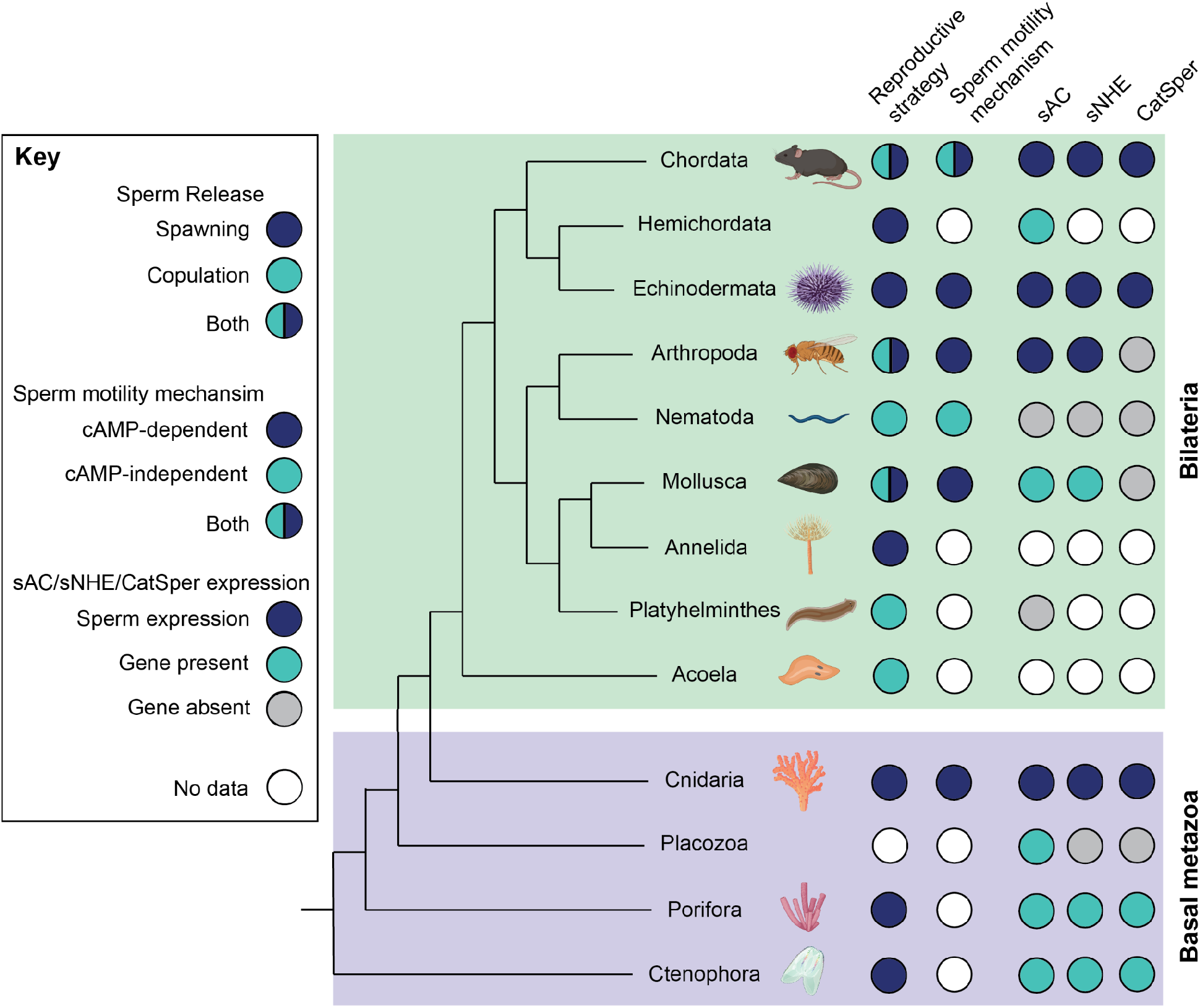
Evolutionary conservation of the sperm motility activation pathway across Metazoan phyla. Left column describes reproductive strategies among the phyla. Spawning refers to release of sperm into an external aquatic environment for internal or external fertilization. Copulation refers to direct sperm transfer and internal fertilization. Middle column indicates phyla where one or more species exhibit cAMP-dependent sperm motility. Right column denotes phyla where one or more species have sAC, SLC9C1 or CatSper expression confirmed in sperm (blue), encoded in the genome (teal), or absent from all available genomes (gray).

## Conclusion

Sexual reproduction is essential for the population growth, evolution, dispersal and community dynamics of marine invertebrates (Crimaldi 2014). While climate change has negatively impacted sexual reproduction across many marine taxa (69), the mechanisms driving these declines remain poorly understood. Ocean warming and acidification, two of the main climate change stressors affecting the ocean, may threaten sperm motility mechanisms that depend on precise intracellular pH by causing cytosolic acidification that may be prohibitively costly to overcome, thus preventing the signaling cascade necessary for motility. Cytosolic acidification may be especially costly to counteract for sperm, which have low cytoplasmic volume, limited energetic resources, and a brief lifespan (62). This could be an important bottleneck for population fitness, as sperm activation and motility are vital for fertilization across metazoan phyla. Importantly, the response of sperm to ocean acidification is nuanced, as both positive and negative shifts in sperm performance have been observed at the level of the individual male in molluscs and echinoderms (70–73). This phenotypic diversity highlights the need to better understand the fundamental mechanisms that regulate sperm performance in order to predict how the fitness of corals and other marine invertebrates will be affected by a changing marine environment.

## Materials and Methods

### Coral sperm intracellular pH motility assays

Egg-sperm bundles were collected from *Montipora capitata* and allowed to break apart naturally in sodium-free seawater (NaFSW; see Supp. Methods for details). Sperm were loaded with 10 µM SNARF-1-AM for 15 min in the dark at room temperature, and fluorescence was imaged using a confocal microscope with 561 nm excitation and dual emission (585 and 640 ± 10 nm) at 25°C. Sperm were imaged before and after addition of 20 mM NH_4_Cl every 20 - 40 seconds over the next 7 min, and at least 14 cells and up to 149 cells were analyzed per time point. Sperm pH_i_ was calculated from the ratio (R) of SNARF1 fluorescence and an in vivo calibration curve generated as previously described (74). Motility was also quantified from the fluorescence images, as non-motile sperm appeared round with smooth edges (Fig. S1A), whereas motile sperm had an oblong shape with an irregular border (Fig. S1B).

### Production of cAMP following activation of motility

Coral sperm suspended in NaFSW were treated with either 20 mM NH_4_Cl or 0.1% DMSO control, and cAMP production was stopped with the addition of HCl (0.17 N final concentration) at 0, 5, 30, 60, 120, and 500 seconds post-exposure. Sperm were lysed by sonication and cAMP content was quantified by ELISA and normalized to total protein.

### Sequence analyses

The presence of key components of the sperm motility activation pathway (sAC, PKA, SLC9C1, CatSper, GC-A, CNGK, and HCN) in corals was investigated by querying two *M. capitata* genomes (38, 43) using echinoderm or cnidarian homologs (see Supp. Methods for more details). Each predicted gene coding sequence was then used as a query in a tBLASTn algorithm search of a sperm-specific RNA-seq database from *M. capitata* sperm (44). *In silico* comparisons of protein structure were carried out using the Clustal Omega multiple sequence alignment tool (75), prediction of transmembrane segments using the TMHMM server (76), and prediction of coiled-coil domains using the DeepCoil server (47).

### Protein expression

Sperm total protein was extracted and protein expression of *mc*sAC and *mc*PKA were detected by Western blotting using custom anti-coral sAC antibodies (37) and commercial anti-PCA antibodies, respectively. Subcellular localization of *mc*sAC was determined by immunocytochemistry. Freshly collected sperm were fixed in 4% paraformaldehyde for 60 min at 4°C, permeabilized in 0.3% triton-X in PBS for 3 min, and incubated with primary antibodies overnight at 4°C. Cells were incubated with secondary antibodies for 1 hour at room temperature in the dark, stained with NucBlue to label nuclei, and imaged using confocal microscopy.

### Phylogenetic analysis of pH-sAC-cAMP signaling within Metazoa

A comparison of the reproductive strategies, sperm motility mechanisms, and sAC/sNHE/CatSper gene conservation was carried out through a search of the literature. Sources used in the compilation of the phylogenetic tree in Figure 4 are described in Figure S5. All gene expression data were derived from (68).

## Acknowledgements

We would like to thank Hollie Putnam, Crawford Drury and the staff at the Hawaii Institute of Marine Biology for logistical support during coral spawning, and Martin Tresguerrues for sharing antibodies. All cartoons were created using BioRender.

## Author contributions

Designed the study: KFS, KLB

Conducted the experiments: KFS, KLB, LAW

Analysed the data: KFS, KLB, LAW, DN

Wrote the paper: KFS, KLB, LAW

