## Supplemental materials for "Molecular mechanisms of sperm motility are conserved in a basal metazoan"

**Speer et al. Supplemental Materials**

### Supplemental Methods

*Coral egg-sperm bundle collections:* Colonies of *Montipora capitata* were housed in flow-through seawater aquaria at the Hawaii Institute of Marine Biology (Special Activity Permit 2020-41). Spawning of *M. capitata* occurs over the five nights following the new moon in the summer months (June-August), starting around 20:45 each evening (Padilla-Gamiño & Gates, 2012), at which time bundles containing sperm and eggs are released from the parent and float to the surface. On each night of collection, bundles were collected from aquaria containing either multiple or isolated individuals. Intact bundles were collected from the surface of the water and placed into sterile plastic containers and immediately brought to the lab for processing.

*Sperm activation and $pH_i$ quantification:* Egg-sperm bundles were allowed to break apart naturally in sodium-free seawater (NaFSW, 430 mM choline chloride, 10 mM calcium chloride, 10 mM potassium chloride, 23 mM magnesium chloride, 25 mM magnesium sulfate, 10 mM HEPES, pH 8.2). Approximately 35 bundles in 3 mL NaFSW were used to attain a sperm concentration of ~1 x $10^7$ cells/$ml^{-1}$. Once the bundles had broken apart, approximately 30-60 min after spawning, the positively buoyant eggs were removed using a transfer pipet. The remaining sperm suspension was diluted 1:4 in NaFSW and loaded with 10 µM SNARF-1-AM in DMSO and 17 mM pluronic acid for 15 min in the dark at room temperature. To quantify resting sperm $pH_i$, 10 uL of the sperm-NaFSW suspension was loaded onto a glass-bottomed dish and imaged using an inverted confocal microscope (Zeiss LSM 710) at 63X magnification (Plan-Apochromat Oil DIC M27 objective, numerical aperture = 1.4). All samples were excited at 561 nm (DPSS, 2% power, 458/561 main beam splitter). SNARF1 fluorescence emission was acquired in two channels simultaneously at 585 and 640 +/- 10 nm, respectively, using a PMT detector (gain = 500). A brightfield image was also captured simultaneously. Images were acquired at 12 bits/pixel using a scan speed of 261.8 Hz. The temperature of the microscope stage was maintained at 25°C.

Basal sperm $pH_i$ was measured over the course of 3 min, after which $NH_4Cl$ was added to a final concentration of 20 mM directly to the sperm suspension and mixed by gently pipetting up and down. Sperm were then imaged every ~20-40 seconds over the next 7 min as described above. A calibration curve relating the ratio (R) of SNARF1 fluorescence to $pH_i$ was generated as previously described (Venn et al. 2009). Briefly, fresh SNARF1-loaded sperm were incubated in calibration solutions of either pH 6.5, 7, 8, or 8.5 in 10 mM nigericin for 5 min and imaged. For comparison, a calibration curve was also generated using adult *M. capitata* cells as previously described (Venn et al. 2009). All images were analyzed in ImageJ by drawing a region of interest (ROI) around each sperm cell and subtracting background fluorescence from an ROI in the surrounding medium from each SNARF1 channel. The fluorescence ratio was then calculated and related to pH using the adult coral calibration curve due to SNARF failing to load in sperm at the pH 7.5 calibration point. At least 14 cells and up to 149 cells were analyzed per time point. Any ROIs saturated at either wavelength were excluded from the analysis. Motility was quantified from these same cells since motile and non-motile sperm were clearly distinguishable: non-motile sperm appeared round with smooth edges (Fig. S1A), whereas motile sperm had an oblong shape with an irregular border (Fig. S1B). Motile cells thin or faint enough to be confused with sperm flagella were excluded.

*Production of cAMP following activation of motility:* Bundles pooled from multiple individuals were allowed to break apart naturally (2.5 ml bundles in 15 ml seawater) over the course of 30-60 min following collection. Eggs were removed from the surface with a pipet, and the remaining sperm suspension was centrifuged at 1500 x g for 5 min. The supernatant was decanted, and the sperm pellet was resuspended in NaFSW. Sperm were added to each of four treatments dissolved in NaFSW: 0.1% DMSO carrier control ± 20 mM $NH_4Cl$ or 50 μM KH7 ± 20 mM $NH_4Cl$. Three replicate assays were run per treatment per time point (10, 30, 60, 120, and 500 seconds) in a volume of 250 μl. Enzyme activity was stopped with the addition of 50 μl of 1N HCl for a final concentration of 0.17 N HCl. Samples were stored at –20° C until quantification of cAMP. Samples were thawed on ice and lysed using a Bioruptor® Pico bath sonicator set to 3 x 1 min sonication cycles (20-60 kHz) with a 30 second cooling period. Lysates were spun down at 1,000 x g for 5 min and cAMP content was measured in supernatants by ELISA (Arbor Assays cAMP EIA Kit, cat. no. K019) using the acetylated format. Experimental samples and standards were diluted in the assay reaction buffer to account for downstream interference by divalent metals (i.e., $Mg^{2+}$). Separate standard curves were run for samples with added $NH_4Cl$. Assay concentrations were calculated using a Four Parameter Logistic Curve fit (https://www.myassays.com/arbor-assays-cyclic-amp-direct-eia-kit-acetyl.assay). cAMP content was normalized to total protein, which was measured from the lysates using the Bradford method with a BSA standard curve (Fisher PI23238).

*Genome & transcriptome mining:* Two draft genomes (Helmkampf et al., 2019, Shumaker et al., 2019) and one RNA-seq database (Van Etten et al., 2020) derived from *M. capitata* sperm were mined to identify transcript-level evidence for the expression of sAC, PKA, SLC9C1, GC-A, CatSper, CNGK, and HCN. To begin, the predicted protein sequence file from an annotated *Montipora capitata* genome database ((Shumaker et al. 2019), http://cyanophora.rutgers.edu/montipora/) was queried using the BLASTp algorithm in the software UGENE to identify gene and predicted amino acid sequences. Either echinoderm (*Strongylocentrotus purpuratus*, *Arbacia punctulata*) or cnidarian (*Euphyllia ancora, Nematostella vectensis)* homologs were used as the query sequence. When an incomplete sequence was identified, a second whole-genome shotgun database from *M. capitata* sperm (Helmkampf et al., 2019) was searched using the tBLASTn algorithm and identical query sequences. The completeness of the predicted *M. capitata* polypeptides varied across genomes but each displayed > 50% similarity to their homolog. The EMBOSS WATER pairwise sequence alignment algorithm was used to calculate the % coverage, Identity and Similarity of *M. capitata* sequences relative to their echinoderm and/or cnidarian homologs. Each predicted gene coding sequence was then used to query an RNA-seq database derived from non-bleached *M. capitata* sperm (Van Etten et al., 2020; BioProject ID: SRX2039373) using a tBLASTn algorithm.

*In silico structural analysis*: The structural characteristics of *M. capitata* sAC, PKA, SLC9C1, GC-A, CatSper, CNGK, and HCN were analyzed by comparing predicted amino acid sequences identified in *M. capitata* genomes to those of metazoan homologs and structurally related proteins. The Clustal Omega multiple sequence alignment tool (Sievers et al. 2011) was used to align *M. capitata* sequences with echinoderm and cnidarian query sequences (Table S2). Mammalian homologs were also included when conservation permitted; specifically for sAC, PKA, SLC9C1, and CatSper. Key amino acid signatures in the *M. capitata* proteins were identified in published structural and functional studies of related proteins. Ion channels with predicted voltage-sensing domains (VSDs), including SLC9C1 and HCN, were also aligned with the *Drosophila melanogaster* Shaker channel whose electromechanical coupling has been extensively studied (Aggarwal and MacKinnon 1996). Protein channels and receptors were searched for hydrophobic transmembrane helices using the TMHMM server (Sonnhammer, von Heijne, and Krogh 1998). The four CatSper subunits were analyzed for C-terminal coiled-coil domains using the DeepCoil server (Ludwiczak et al. 2019).

*Protein expression:* Bundles from three separate colonies were allowed to break apart naturally in seawater and eggs were removed. Sperm were pelleted by centrifugation (1500 x g for 5 min) and resuspended in 100 mM Tris pH 7.5, 150 mM NaCl, 100 uM EDTA with protease inhibitors (Sigma P8340) and stored at –80° C. Sperm were thawed on ice and lysed using a Fisher Homogenizer 850 (10 sec at 25,000 rpm). Samples were spun down at 1,000 x g for 5 min to remove cellular debris, supernatants were collected, and total protein concentration was measured as described above. Next, 7.5 µg of total protein was loaded onto a Novex 4-12% Tris-Glycine Mini Gel and transferred in Towbin's buffer without alcohol for 100 min at 100V. Immunological detection of *M. capitata* sAC (mcsAC) was performed using custom affinity-purified polyclonal anti-coral sAC antibodies (GenScript USA, Inc., 0.55ug/mL) raised against an antigen peptide in the C2 catalytic region of sAC from the coral *Pocillopora damicornis* (LPGDKHEDDPARAL; (Barott, Barron, and Tresguerres 2017)). This sequence is identical to the predicted *m*csAC protein sequence from the *M. capitata* genome (Shumaker et al. 2019). Specificity of these antibodies for *mc*sAC was confirmed by Western blot using a preincubation of 300-fold molar excess of the immunizing peptide as a control. The PKA western was conducted using commercial antibodies (Cell Signaling #4781, 1:1000) recognizing the C-terminus of the alpha catalytic subunit of human PKA (PKA C$\alpha$; 82.1% identity to *mc*PKA C$\alpha$; Table S1). Commercial antibodies were used as loading controls, including: Actin (Developmental Studies Hybridoma Bank #JLA-20, 0.3 ug/mL) and β-tubulin (Developmental Studies Hybridoma Bank #E7, 0.5 ug/mL).

*Immunolocalization:* Freshly collected sperm were fixed in 4% paraformaldehyde and allowed to attach to glass slides for 60 min at 4°C. Sperm were rinsed in cold PBS and permeabilized in 0.3% triton-X in PBS for 3 min at room temperature. For a subset of samples, fresh sperm were allowed to attach to silane coated glass slides for 30 min at room temperature, and then fixed in 4% PFA in S22 buffer for 10 min at room temperature. All samples were then incubated in blocking buffer (1% BSA and 0.01% keyhole limpet hemocyanin in PBS) for 30 min at 4°C. Blocking buffer was removed and samples were incubated with primary antibodies overnight at 4°C. All antibodies were diluted in blocking buffer to the following working concentrations: anti-coral sAC, 5.5 µg/ml; anti-β-tubulin, 3.5 µg/ml; goat anti-rabbit AlexaFluor 488 (Invitrogen A32731), 4 µg/ml; goat anti-mouse AlexaFluor 594 (Invitrogen A32742), 4 µg/ml. Primary antibodies were removed with a pipet and slides were rinsed by immersion in cold PBS three times for 10 seconds each. Secondary antibodies were added and samples were incubated for 1 hour at room temperature in the dark. Slides were again rinsed in cold PBS, mounted using

Prolong Glass Antifade with NucBlue (Invitrogen P36983), and cured overnight in the dark at room temperature. All slides were then stored at -20°C in the dark. Sperm were imaged using a confocal microscope (Leica SP8).

### Supplemental Tables

Table S1. Genomic and transcriptomic identification of sperm motility pathway.

| Protein Name | *M. capitata* Genome ID | Transcripts?[3] (no. reads >95% identical) | Homolog [NCBI ID] | Homolog Coverage Length (aa) | Identity/ Similarity (%) |
|---|---|---|---|---|---|
| *mc*sAC | adi2mcaRNA12271_R0[1] | Y (68) | *S. purpuratus* [Q4VGX1] | 76 - 1864 (1868) | 33.2/48.3 |
| | | | *P. damicornis* [APM87498.1] | 24 - 1889 (1891) | 65.7/75.7 |
| *mc*PKA (Cα) | augustus.g2600.t1[1] | Y (100) | *S. purpuratus* [XP_030852244.1] | 1 – 352 (352) | 79.5/90.6 |
| *mc*SLC9C1 | augustus.g27233.t1[1] *and* augustus.g27234.t1[1] | Y (9) | *S. purpuratus* [NP_001091927] | 130 - 1064 (1325) | 52.5/69.8 |
| *mc*GC-A | RDEB01000014.1[2] | Y (29) | *Euphyllia ancora* [MH894389] | 14 - 1049 (1049) | 67.6/78.0 |
| | | | *S. purpuratus* [P16065] | 44 - 1044 (1125) | 28.0/42.3 |
| *mc*CatSper (α1) | augustus.g52025.t1 | N (0) | *S. purpuratus* [XP_030854941.1] | 285 - 410 (474) | 50.8/68.8 |
| *mc*CatSper (α2) | augustus.g3319.t1[1] | Y (2) | *S. purpuratus* [XP_030832241.1] | 18 - 392 (517) | 51.1/70.7 |
| *mc*CatSper (α3) | augustus.g52671.t1[1] | Y (1) | *S. purpuratus* [XP_030835949.1] | 57 - 390 (394) | 38.6/51.5 |
| *mc*CatSper (α4) | adi2mcaRNA3204_R1[1] | Y (9) | *S. purpuratus* [XP_030846926.1] | 3 – 223 (547) | 53.4/70.2 |
| *mc*CNGK | adi2mcaRNA7057_R0[1] *and* adi2mcaRNA7058_R0[1] | Y (69) | *Arbacia punctulata* [MT557712.1] | 33 - 2069 (2242) | 38.9/53.9 |
| *mc*HCN | RDEB01000437.1[2] | Y (10) | *S. purpuratus* [NP_999729.1] | 179 – 666 (767) | 52.3/73.1 |

Nucleotide sequences obtained from: [1(Shumaker et al. 2019)], [2(Helmkampf et al. 2019)], [3(Van Etten et al. 2020)].

Table S2. Homolog sequences aligned with *M. capitata* proteins using the Clustal Omega server.

| **Protein name** | **MSA sequences [NCBI ID]** |
|---|---|
| *mc*sAC | *S. purpuratus* [Q4VGX1]<br>*P. damicornis* [APM87498.1]<br>*M. musculus* [Q8C0T9] |
| *mc*PKA (Cα) | *S. purpuratus* [XP_030852244.1]<br>*H. sapien* [P17612] |
| *mc*SLC9C1 | *S. purpuratus* [NP_001091927]<br>*M. musculus* [AAQ88278.1]<br>*D. melanogaster* Shaker [CAA29917.1] |
| *mc*GC-A | *E. ancora* [[MH894389]<br>*S. purpuratus* [P16065] |
| *mc*CatSper (α1) | *N. vectensis* [XP_001641973.1]<br>*S. purpuratus* [XP_030854941.1]<br>*M. musculus* [Q91ZR5] |
| *mc*CatSper (α2) | *N. vectensis* [XP_001629309.1]<br>*S. purpuratus* [XP_030832241.1]<br>*M. musculus* [A2ARP9] |
| *mc*CatSper (α3) | *N. vectensis* [XP_001629965.1]<br>*S. purpuratus* [XP_030835949.1]<br>*M. musculus* [Q80W99] |
| *mc*CatSper (α4) | *N. vectensis* [XP_001631669.1]<br>*S. purpuratus* [XP_030846926.1]<br>*M. musculus* [Q8BVN3] |
| *mc*CNGK | *A. punctulata* [QKM75728.1]<br>*P. damacornis* [XP_027036539.1] |
| *mc*HCN | *S. purpuratus* [NP_999729.1] |

Table S3. Transmembrane helix predictions carried out using the TMHMM server (Sonnhammer, von Heijne, and Krogh 1998).

| Protein Name | Protein ID | TMHMM Analysis/Predicted based on homology |
|---|---|---|
| *mc*SLC9C1 | augustus.g27233.t1[1] *and* augustus.g27234.t1[1] | 13/18[a] [10 NHE & 3 VSD] |
| *pd*SLC9C1 | XP_027042233.1[3] | 15/18[a] [12 NHE & 3 VSD] |
| *mc*GC-A | RDEB01000014.1[2] | 0/1[b] |
| *mc*CatSper(α1) | augustus.g52025.t1[1] | 5/6[a] |
| *mc*CatSper(α2) | augustus.g3319.t1[1] | 3/6[a] |
| *mc*CatSper(α3) | augustus.g52671.t1[1] | 2/6[a] |
| *mc*CatSper(α4) | adi2mcaRNA3204_R1[1] | 5/6[a] |
| *mc*CNGK | adi2mcaRNA7057_R0[1] *and* adi2mcaRNA7058_R0[1] | 18/24[c] [4 R1, 6 R2, 4 R3, 4 R4] |
| *mc*HCN | RDEB01000437.1[2] | 3/6[a] |

[1]Shumaker et al., 2019, [2]Helmkampf et al., 2019, [3]NCBI Protein database, [a]Compared to *Strongylocentrotus purpuratus* homolog, [b]Compared to *Euphyllia ancora* homolog, [c]Compared to *Arabica punctulata* homolog

Table S4. Coiled-coil domain predictions in CatSper carried out using the DeepCoil server (Ludwiczak et al. 2019).

| Protein Name | C-terminal coiled-coil probability >0.5? |
|---|---|
| *mc*CatSper(α1) | N |
| *mc*CatSper(α2) | Y |
| *mc*CatSper(α3) | Y |
| *mc*CatSper(α4) | N |

Table S5. Data sourced for the comparison of reproductive strategies, sperm motility mechanisms, and sAC/sNHE/CatSper gene expression in Figure 4.

| Taxa | Reproductive Strategy | Sperm Motility Mechanism | sAC (sperm expression) | sNHE (sperm expression) | CatSper (sperm expression) |
|---|---|---|---|---|---|
| **Chordata** | (Extavour 2007) | (Kaupp, Hildebrand, and Weyand 2006) | (Buck et al. 1999) | (Wang et al. 2003) | (Quill et al. 2001; Ren et al. 2001) |
| **Hemichordata** | (Extavour 2007) | | | | |
| **Echinodermata** | (Extavour 2007) | (Kaupp, Hildebrand, and Weyand 2006) | (Bookbinder, Moy, and Vacquier 1990) | (Windler et al. 2018) | (Seifert et al. 2015) |
| **Arthropoda** | (Extavour 2007) | (Osanai, Kasuga, and Aigaki 1989) | (Romero and Nishigaki 2019) | (Romero and Nishigaki 2019) | |
| **Nematoda** | (Extavour 2007) | (Smith 2018) | | | |
| **Mollusca** | (Extavour 2007) | (Boulais et al. 2019) | | | |
| **Annelida** | (Fischer 1999) | | | | |
| **Platyhelminthes** | (Rawlinson et al. 2008) | | | | |
| **Acoela** | (Achatz et al. 2013) | | | | |
| **Cnidaria** | (Extavour 2007) | **Fig. 1** | **Fig. 2** | **Fig. 3** | **Fig. 3** |
| **Placozoa** | (Extavour 2007) | | | | |
| **Porifora** | (Extavour 2007) | | | | |
| **Ctenophora** | (Extavour 2007) | | | | |

### Supplemental Figures

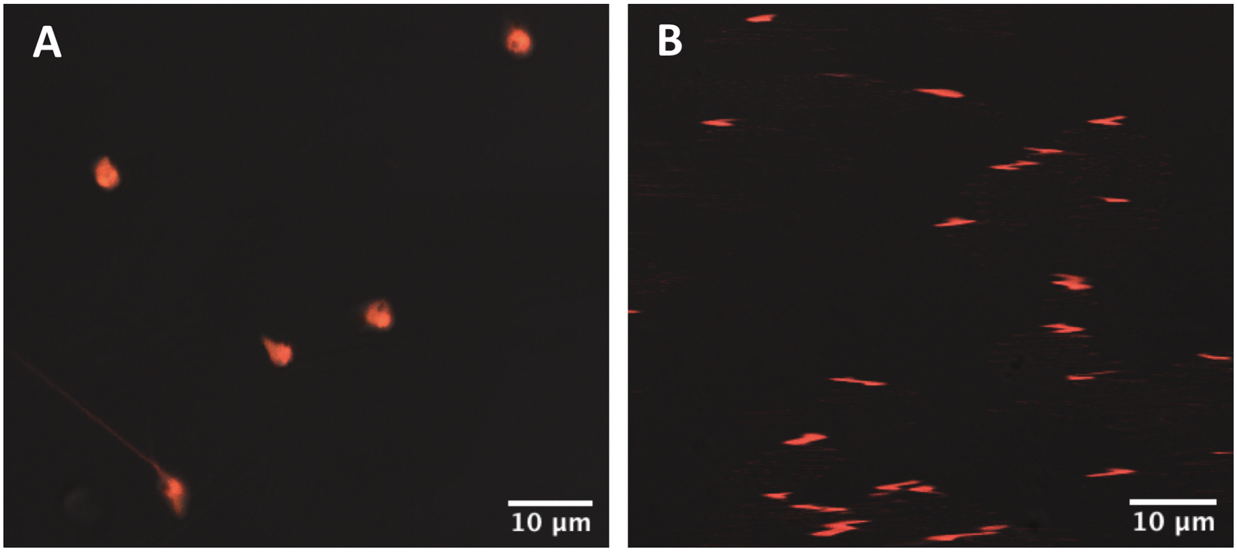


Figure S1. Representative images of SNARF1-AM loaded *Montipora capitata* sperm A) before $NH_4Cl$ addition (t = 0 s) and B) after $NH_4Cl$ addition (t = 45 s).

```
Mcap_sAC        DD-------------  503
Pdam_sAC        DREE-------EEEE  515
Mmus_sAC        EKVMFGMAYLICNRY  481
```

Figure S2. Conservation of an autoinhibitory peptide in soluble adenylyl cyclase (sAC). The minimum autoinhibitory region of *M. musculus* sAC (Mmus_sAC) is defined by the peptide sequence ((Chaloupka et al. 2006)) that separates the longest sAC isoform with high catalytic activity from the shortest sAC isoform that is inhibited (Chaloupka et al. 2006). The sequence is missing in both *M. capitata* sAC (Mcap_sAC) and *P. damicornis* sAC (Pdam_sAC).

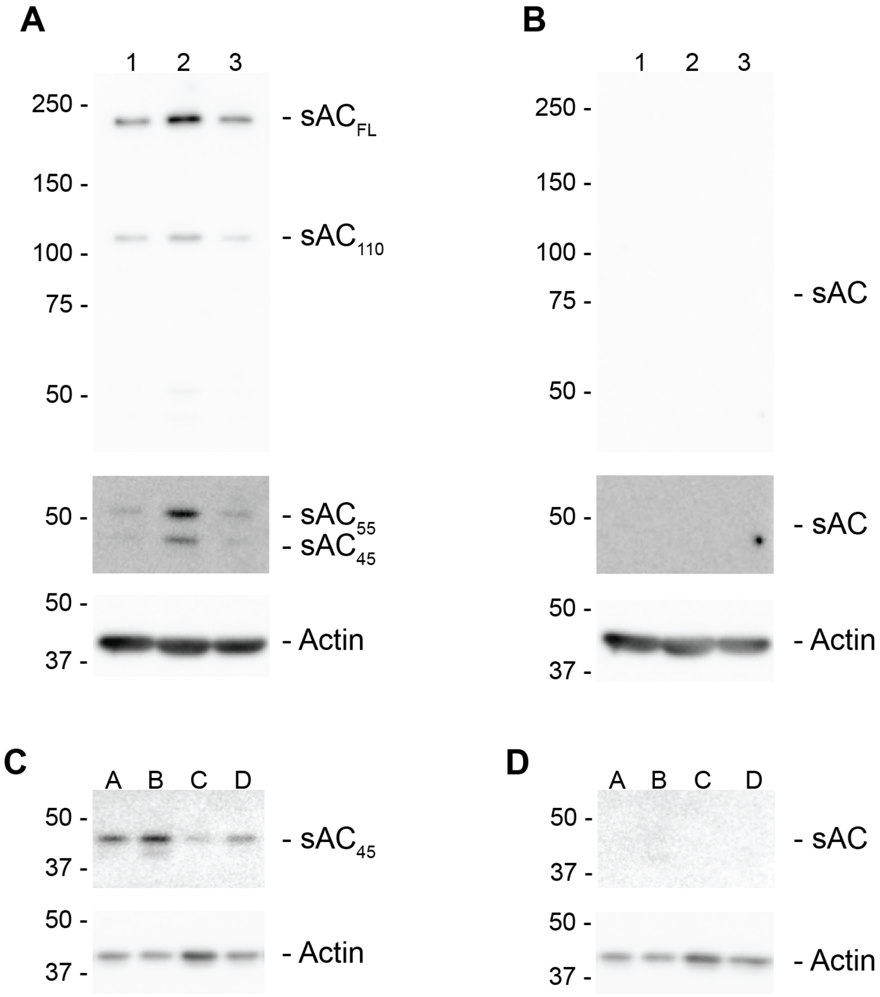


Figure S3. sAC antibody is highly specific and detects multiple isoforms of the enzyme. Western blots performed on *Montipora capitata* (A-B) sperm from three individuals (1, 2 & 3); (C-D) four adults (A, B, C & D). Blots preincubated with antigen peptide on the right (B,D).

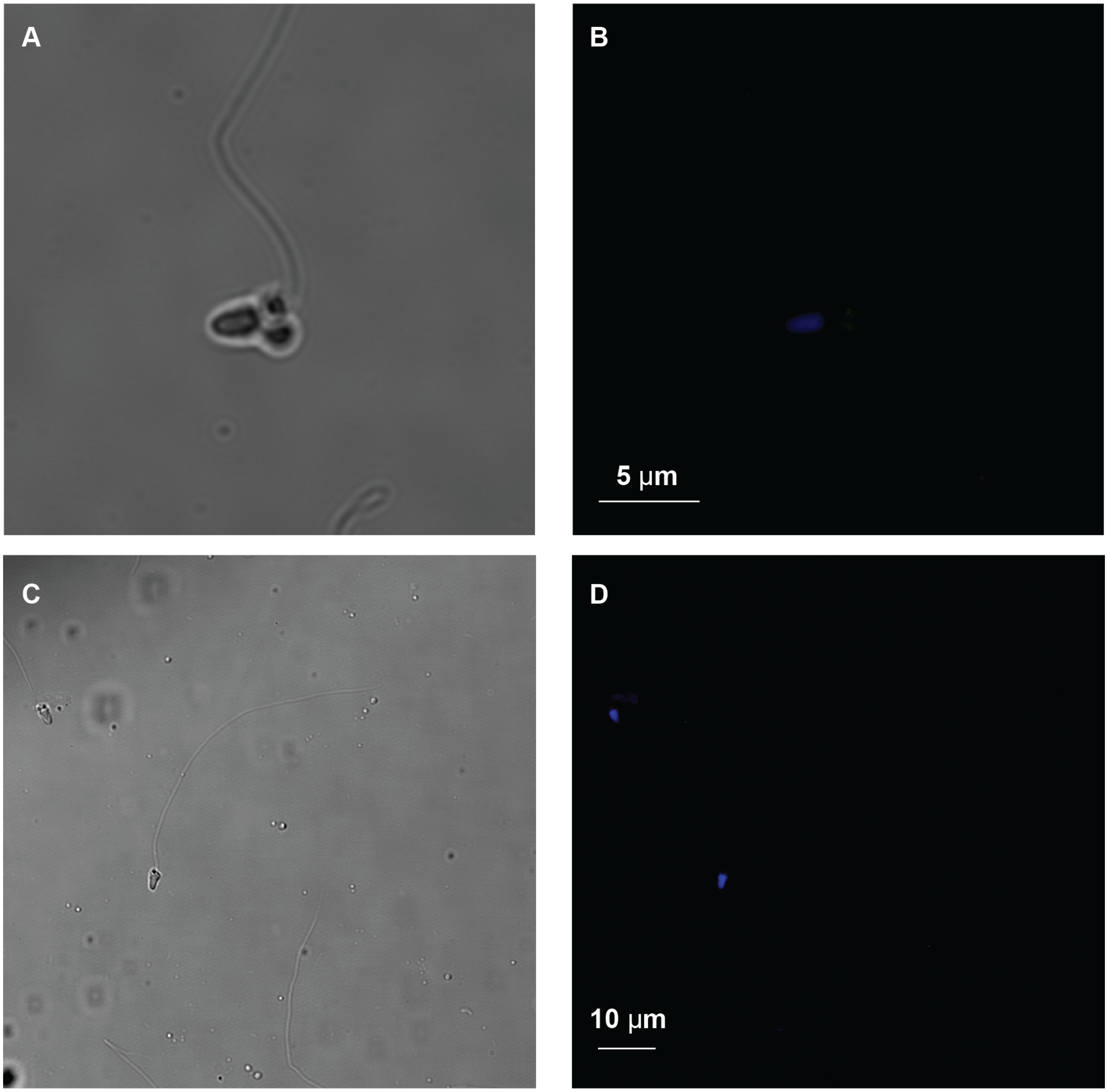


Figure S4. Controls for immunofluorescent staining of *M. capitata* sperm with secondary antibodies only (no primary). A) Brightfield and B) fluorescent signals captured for sperm stained with goat anti-rabbit AlexaFluor 488, goat anti-mouse 594, and NucBlue imaged at 7X Zoom; identical to Fig. 2E-G. C) Brightfield and D) fluorescent signals for sperm fixed on silane-treated slides & stained with goat anti-rabbit AlexaFluor 488 and NucBlue imaged at 2X Zoom; identical to Fig. 2H, I.

**A**

```
                         NHE S3                                   NHE S4
Mcap_SLC9C1    WTWFEALLFGSIVSATDPVAVVALLNDLGTSKQLSTIIEGESLLNDGMAIVLYKIFFNLAFSS  127
Spur_SLC9C1    WNFSEAMMFGAIMSATDPVAVVALLKDLGASKQLGTIIEGESLLNDGCAIVIFNVFMKMVFFP  255
Mmus_SLC9C1    -----WLLFSAVLISSDPMLTSASIRDLGLSRSLTNLINGESLLTSVLSLVIYSGVVHIRFKS  163
```

**B**

```
                    VSD S4
Mcap_SLC9C1    PSILKVAKVFRVLRMGRVLRLFKTLL  652
Spur_SLC9C1    LSSIKVVKLFRLLRGLRMLRLTKALI  818
Mmus_SLC9C1    FDLTETVVFMNVIRLLRILRILKLVT  692
Dmel_Shaker    LAILRVIRLVRVFRIFKLSRHSKGLQ  353
```

Figure S5. Sequence comparison of *Montipora capitata* SLC9C1 domains with bilaterian homologs. A) Alignment of the cation binding motif of the SLC9C1 family. *M. capitata* sequence identified from (Shumaker et al. 2019) (Mcap_SLC9C1), *S. purpuratus* SLC9C1 (Spur_SLC9C1), *M. musculus* (Mmus_SLC9C1). The conserved motif (bold, blue) was identified within the second and third transmembrane domains (S3 & S4) of the NHE domain. B) Alignment of SLC9C1 family voltage-sensing domains (VSD) with the well-studied VSD of the *D. melanogaster* Shaker potassium channel (Dmel_Shaker). Positively charged amino acids (bold, blue), spaced at a three amino acid interval, are responsible for electromechanical coupling.

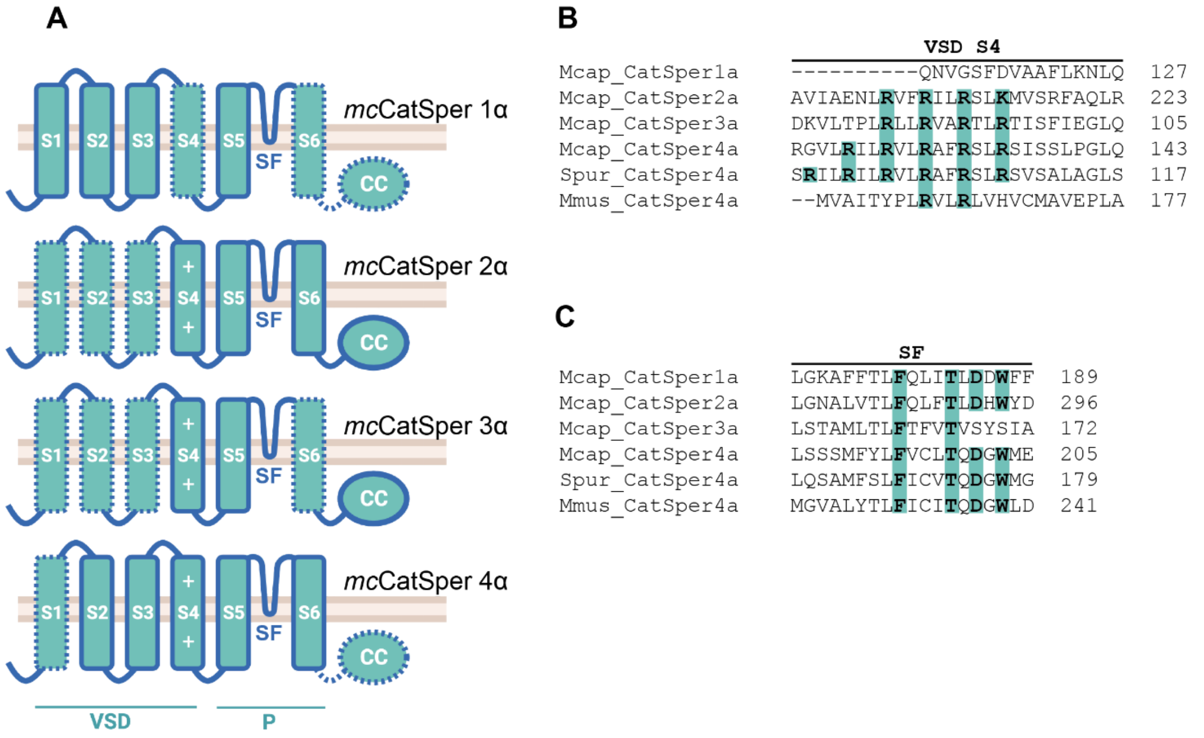


Figure S6. Identification and comparison *Montipora capitata* CatSper peptides 1-4α with bilaterian homologs. A) Domain structure of *mc*CatSper peptides 1-4α identified in an *M. capitata* genome (Shumaker et al. 2019). Alignment of *mc*CatSper peptides with *S. purpuratus* homologs identified shared sequence (solid lines) and sequence missing from *M. capitata* genes (dotted lines). Transmembrane segments S1-4 form a voltage-sensing domain (VSD) with a series of positively charged amino acids (+) in S4 responsible for electromechanical coupling. Transmembrane segments S5, 6 and their intermediary selectivity filter (SF) create the channel pore (P). A C-terminal coiled-coil (CC) domain is responsible for oligomerization. B) *mc*CatSper peptides 2, 3 and 4α contained 4-5 positively charged residues in VSD S4. Bilaterian CatSper peptides vary significantly in the number of S4 charges; *A. punctulata* CatSper4α contains six and *M. musculus* CatSper4α contains two. C) *mc*CatSper peptides 1, 2, and 4α contained the canonical [T/S]x[D/E]xW motif of $Ca^{2+}$-selective channels in their selectivity filter, similar to *S. purpuratus* CatSper peptide 4α (Spur_CatSper4a) and *M. musculus* CatSper peptide 4α (Mmus_CatSper4a).

```
                    SF
Mcap_CNGK_dom1   IYWATATATGTGYGDIHAV  254
Mcap_CNGK_dom2   LYWAAATMSSTGYGDIHAH  700
Mcap_CNGK_dom3   LYWAAATSASVGYGDIHAH  1262
Mcap_CNGK_dom4   AYWVVATMTSTGYGDIHAH  1779
Apun_CNGK_dom4   IYWAVATLTSTGYGDIHAY  1994
Dmel_Shaker      FWWAVVTMTTVGYGDMTPV  421
```

Figure S7. The selectivity filters of the four *Montipora capitata* CNGK subunits (Mcap_CNGK_dom1-4) contain the T/S rich region and GYG motif of $K^{+-}$-selective channels (bold, blue) also found in *A. punctulata* CNGK domain 4 (Apun_CNGK_dom4) and the canonical *D. melanogaster* $K^{+}$-selective Shaker channel (Dmel_Shaker).

**A**

```
                     SF
Mcap_HCN    YSWALFKAMSHMLCIGYGRYPPQA  256
Spur_HCN    YTWALFKALSHMLCIGYGKFPPQS  438
```

**B**

```
                   VSD  S4
Mcap_HCN    LRILKLAKLLSLLRLLRVSRLMRYIT  174
Spur_HCN    LKILRFAKLLSLLRLLRLSRLMRFVS  352
Dmel_Shaker LAILRVIRLVRVFRIFKLSRHSKGLQ  353
```

Figure S8. Key functional domains of the *Montipora capitata* HCN (Mcap_HCN) channel include A) the selectivity filter (SF), which contains a GYG motif (bold, blue) of cation-binding channels like its *S. purpuratus* (Spur_HCN) homolog and B) the voltage-sensing domain (VSD), which contains seven positively charged residues embedded in transmembrane segment S4. Unlike the *D. melanogaster* Shaker channel (Dmel_Shaker), these residues are split into two clusters of 3-4 residues.

Romero, Francisco, and Takuya Nishigaki. 2019. "Comparative Genomic Analysis Suggests That the Sperm-Specific Sodium/proton Exchanger and Soluble Adenylyl Cyclase Are Key Regulators of CatSper among the Metazoa." *Zoological Letters* 5 (July): 25.

Seifert, Reinhard, Melanie Flick, Wolfgang Bönigk, Luis Alvarez, Christian Trötschel, Ansgar

Poetsch, Astrid Müller, et al. 2015. “The CatSper Channel Controls Chemosensation in Sea Urchin Sperm.” *The EMBO Journal* 34 (3): 379–92.
Shumaker, Alexander, Hollie M. Putnam, Huan Qiu, Dana C. Price, Ehud Zelzion, Arye Harel, Nicole E. Wagner, Ruth D. Gates, Hwan Su Yoon, and Debashish Bhattacharya. 2019. “Genome Analysis of the Rice Coral Montipora Capitata.” *Scientific Reports* 9 (1): 2571.
Sievers, Fabian, Andreas Wilm, David Dineen, Toby J. Gibson, Kevin Karplus, Weizhong Li, Rodrigo Lopez, et al. 2011. “Fast, Scalable Generation of High-Quality Protein Multiple Sequence Alignments Using Clustal Omega.” *Molecular Systems Biology* 7 (1): 539.
Smith, Harold E. 2018. *Nematode Sperm Motility*. WormBook.
Sonnhammer, E. L., G. von Heijne, and A. Krogh. 1998. “A Hidden Markov Model for Predicting Transmembrane Helices in Protein Sequences.” *Proceedings / ... International Conference on Intelligent Systems for Molecular Biology ; ISMB. International Conference on Intelligent Systems for Molecular Biology* 6: 175–82.
Van Etten, Julia, Alexander Shumaker, Tali Mass, Hollie M. Putnam, and Debashish Bhattacharya. 2020. “Transcriptome Analysis Provides a Blueprint of Coral Egg and Sperm Functions.” *PeerJ* 8 (August): e9739.
Venn, A. A., E. Tambutté, S. Lotto, D. Zoccola, D. Allemand, and S. Tambutté. 2009. “Imaging Intracellular pH in a Reef Coral and Symbiotic Anemone.” *Proceedings of the National Academy of Sciences* 106 (39): 16574–79.
Wang, Dan, Shelby M. King, Timothy A. Quill, Lynda K. Doolittle, and David L. Garbers. 2003. “A New Sperm-Specific Na+/H+ Exchanger Required for Sperm Motility and Fertility.” *Nature Cell Biology* 5 (12): 1117–22.
Windler, F., W. Bönigk, H. G. Körschen, E. Grahn, T. Strünker, R. Seifert, and U. B. Kaupp. 2018. “The Solute Carrier SLC9C1 Is a Na /H -Exchanger Gated by an S4-Type Voltage-Sensor and Cyclic-Nucleotide Binding.” *Nature Communications*. https://doi.org/10.1038/s41467-018-05253-x.
